# Aberrant cerebrovascular reactivity presents as an early biomarker of psychosis susceptibility in patients with 22q11.2DS

**DOI:** 10.1101/2025.06.02.657392

**Authors:** Farnaz Delavari, Stefano Moia, Silas Forrer, Corrado Sandini, Nada Kojovic, Alessandro Pascucci, Clémence Lacour, Stephan Eliez, Dimitri Van De Ville

## Abstract

The brain’s ability to regulate blood flow is fundamental to both its function and development. In the context of neurodevelopmental disorders such as schizophrenia, understanding the complex interactions between cerebrovascular health and brain function is crucial for unraveling the pathophysiology of psychosis. This study investigates the developmental trajectory of cerebrovascular reactivity (CVR) in 22q11 deletion syndrome (22q11.2DS) compared to healthy controls, and its association with psychosis susceptibility. Using a longitudinal data-set of resting-state fMRI, we mapped voxel-level CVR across development. We found significant and early CVR impairments in 22q11.2DS, and in particular in those who later developed positive psychotic symptoms (PPS+). These impairments were evident within the anterior cingulate cortex, frontal lobes, and globi pallidi (GB). We propose that the pattern of CVR reduction presenting early during childhood is possibly linked to blood brain barrier impairment. A decrease in CVR during childhood and within the frontal regions and GB was predictive of subsequent development of positive psychotic symptoms (PPS), which often occurs during adolescence in 22q11.2DS patients. These findings suggest that cerebrovascular health is critical for normal brain development, particularly in regions like the striatum, which are vulnerable to vascular damage due to their anatomical features. These results underline the potential of CVR as an early biomarker for psychosis vulnerability, emphasizing the need for targeted interventions to mitigate neurodevelopmental disruptions of cerebrovascular health in 22q11.2DS.

## Introduction

Cerebrovascular reactivity (CVR), defined as the capacity of cerebral blood vessels to adjust their diameter in response to vasodilatory stimuli, such as increased CO2 concentration in the bloodstream, plays a pivotal role in maintaining gas homeostasis and cerebral perfusion and interacts with neurovascular coupling (Fisher and Mikulis 2021). This complex physiological phenomenon, involving endothelial cells, pericytes, astrocytes, and neurons, plays a crucial role in maintaining brain function and responsiveness to metabolic demands (Hall, Reynell et al. 2014, Howarth, Sutherland et al. 2017). The coupling between vessels and neurons is mostly studied in the neuro-to-vascular direction. However, the vascular tone has also a direct impact on neuronal activity in the cortex (Kim, Ramiro Diaz et al. 2016). Thus, the clinical implications of CVR are broad and impact various conditions of the brain. Indeed, CVR has been well studied in the domain of aging and cognitive function, where it has been shown that a decline in CVR predicts deterioration in cognitive functioning in older adults(Peng, Chen et al. 2018).

The implications of CVR abnormalities extend to neurodegenerative diseases, such as Alzheimer’s and Parkinson’s disease (Ryman, Shaff et al. 2023,(Sur, Lin et al. 2020), where decreased CVR is discussed to result in chronic hypoxia, giving rise to neuroinflammation and processes that contribute to disease progression. In the case of neuroinflammatory disorder, for instance multiple sclerosis (MS), chronic inflammation might lead to a reduced CVR, contributing to symptoms such as cognitive dysfunction and fatigue (D’Haeseleer, Hostenbach et al. 2015, Pelizzari, Laganà et al. 2020). Aside from global CVR reduction, regional variabilities in cerebrovascular reactivity, such as those observed particularly in the hippocampus, frontal, cingulate, and parietal areas, contribute to varied cognitive outcomes with age (Catchlove, Parrish et al. 2018, McKetton, Sobczyk et al. 2018). These findings underscore the importance of spatial heterogeneity in vascular decline that may give rise to a unique set of neurological symptoms.

Studies investigating CVR in psychiatric disorders are scarce. Impaired perfusion has been associated with anxiety and other mood disorders (Harandi, Kimia et al. 2023). Regions within the brainstem and amygdala expressing altered CVR have been implicated in mice models of anxiety disorders (Wenzel, Hansen et al. 2020). Moreover, a study of bipolar disorder among adolescents documented reduced CVR, emphasizing on the importance of considering developmental hits to the cerebrovascular health in psychiatric disorders (Urback, Metcalfe et al. 2019). Of note, to our knowledge there are no studies on CVR in children and adolescent populations in the context of prodromal phase of psychotic disorders, despite the evidence demonstrating that CVR measurements are reproducible in children. For instance, studies in children with risk of ischemia have demonstrated that CVR measurements can indicate tissue-level microvascular dysfunction (Kim, Leung et al. 2016, Leung, Kim et al. 2016, Dlamini, Yau et al. 2017), highlighting the utility of CVR assessments in pediatric studies. Indeed, it is imaginable for neurodevelopmental processes to rely heavily on cerebrovascular dynamics. CVR disruption during critical periods of brain development may lead to aberrant structural and functional alterations, consequently contributing to the emergence of psychopathological manifestations later in life (Kim, Leung et al. 2016). Understanding how cerebrovascular dynamics affects abnormal brain development and psychopathology holds a significant promise for revealing underlying mechanisms and developing targeted interventions aimed at preventing neurodevelopmental disorders.

Schizophrenia, viewed primarily as a disorder characterized by onset of psychosis, is now increasingly recognized as a neurodevelopmental disorder with origins in early brain development (Insel 2010). This view highlights the importance of investigating the developmental trajectories preceding the onset of first symptoms of psychosis. However, characterizing brain maturation during the early premorbid stages of schizophrenia presents significant challenges, primarily due to the low incidence of psychosis in the general population (Charlson, Ferrari et al. 2018). To address this challenge, individuals with 22q11.2DS present a particularly valuable model (Gur, Roalf et al. 2021). With a 40% risk of progressing into psychosis, 22q11.2DS provides a unique opportunity to investigate the neurodevelopmental abnormalities that give rise to the onset of schizophrenia (Schneider, Schaer et al. 2014). Additionally, 22q11.2DS represent a unique population for investigating the interplay between genetic predispositions, cerebrovascular dysfunction, and psychiatric manifestations.

Various mechanisms in 22q11.2DS may affect vascular health and thus cerebrovascular function, including endothelial dysfunction and vascular malformations that lead to an unstable vascular network, potentially impairing blood supply to the brain (Yi, Tang et al. 2014, Unolt, Versacci et al. 2018). Moreover, the deletion area encompasses the gene responsible for maintaining the integrity of the blood-brain barrier (BBB), impaired expression of which has been associated with presentation of psychosis (Greene, Kealy et al. 2018, Crockett, Kebir et al. 2024). Moreover, a recent study demonstrated early impairment of the BBB in the endothelium of the brains of mice-models with gene-deletion equivalent to 22q11.2DS (Crockett, Ryan et al. 2021). It is worth noting, a study using focused ultrasound on rat brains demonstrated that the regional opening of the BBB results in attenuated blood oxygenation level dependent (BOLD) signal in the affected region (Todd, Zhang et al. 2019). Furthermore, they demonstrated that a portion of this attenuation is attributed to a vascular component, as evidenced by impaired CVR during a gas challenge in this animal model. Thus, BBB impairment results in decreased CVR which is measurable by the BOLD signal. These findings are critical as the BOLD measurement could be translated into human research for the assessment of BBB integrity.

Performing CVR mapping on children is feasible (Leung, Kim et al. 2016) but challenging nonetheless. Indeed, gold standard approaches such as acetazolamide injection or gas challenges (Fisher and Mikulis 2021) are either not indicated or can cause discomfort in vulnerable subjects, such as children. Breathing challenges are normally employed as a more approachable and practical alternative, and while they are robust to certain lack of compliance (Solis-Barquero, Echeverria-Chasco et al. 2021), they are still too challenging for pediatric populations. However, a recent advancement in non-invasive functional magnetic resonance imaging (MRI) technique allows for the qualitative assessment of resting-state CVR (rCVR) (Liu, Li et al. 2017), removing completely the constraint of subject compliance and providing an opportunity to employ this measurement in a developing human population to gain insights into neurovascular coupling and cerebrovascular health.

In this study, we leverage the large cohort of Geneva-22q11.2DS who underwent longitudinal resting state BOLD functional MRI, to measure rCVR, hereafter called CVR, in individuals with 22q11.2DS and healthy controls, aiming to characterize developmental changes in cerebrovascular function across the whole brain. Our investigation focuses on identifying if CVR impairments are present in patients with 22q11.2DS. Moreover, we aim to show when during the course of the development these impairments present themselves and whether they may contribute to the emergence of positive psychotic symptoms. Furthermore, we aim to investigate if part of these changes could be attributed to the congenital heart defects present in this population at birth. Given the strong genetic component on vascular system development and integrity (Cioffi, Martucciello et al. 2014, Greene, Hanley et al. 2019), individuals with 22q11.2DS possess inherent vulnerabilities in cerebrovascular health, which may play a role in their predisposition to a spectrum of neurodevelopmental abnormalities. We hypothesize that the markers of CVR impairment in critical brain regions are present at an early age already during childhood. We further hypothesize that cerebrovascular alteration are an early hit that are associated with later development of psychosis.

## Material and Methods

### Participants

A total of 113 participants diagnosed with 22q11.2DS (49% female, age range 6-30 years) – with a confirmed genetic diagnosis of 22q11.2 deletion syndrome from a local genetics department, and 133 healthy control subjects (HCs) (56% female, age range 6–30 years) underwent multiple MRI scans. For the sample of HCs, we included participants from unaffected siblings of the patients or through an open call to the Geneva State School system in Switzerland. HCs were screened for history of any psychopathology and developmental abnormalities or cardiac anomalies and were matched for age and sex.

Deletion carriers underwent a comprehensive clinical assessment with an expert psychiatrist (SE) during each visit, during which Structured Interview for Prodromal Syndromes(Miller, McGlashan et al. 2003) (SIPS) was administered to deletion carriers to assess the presence of attenuated symptoms of psychosis. One participant was too young and was not assessed with SIPS. As studied in previous publications from the same cohort (Scariati, Schaer et al. 2014, Zöller, Padula et al. 2018), we used SIPS results to create subgroups based on the participants’ clinical characteristics with regards to psychosis vulnerability. Patients were classified as PPS(+) (n = 46) if they scored equal to or higher than 3 in at least one visit on any of the subscales for positive psychotic symptoms (Unusual Thought Content, Suspiciousness, Grandiosity, Hallucinations, and Disorganized Communication) (Fusar-Poli, Borgwardt et al. 2013). The remaining participants were considered nonPPS. Participants’ clinical characteristics are further detailed in Table 1.

**Table 1.** Table of demographics and participant characteristics IQ measurement: Wechsler Intelligence Scale for Children-III for children(Wechsler and Kodama 1949) and Wechsler Adult Intelligence Scale-III for adults (Wechsler 1955). Presence of psychiatric disorders: Clinical interview with patients using the Diagnostic Interview for Children and Adolescents–Revised(Reich 2000), the psychosis supplement from the Schedule for Affective Disorders and Schizophrenia for School-Age Children–Present and Lifetime version(Kaufman, Birmaher et al. 1997), and the Structured Clinical Interview for DSM-5 Axis I Disorders(First 1997).NaN, not available; PPS(+), with positive psychotic symptoms; nonPPS, without positive psychotic symptoms; 22q11.2DS, 22q11 deletion syndrome.

|  | **22q11.2DS** | **Healthy Control** | **P-value** | **PPS+** | **nonPPS** | **P-value** |
| --- | --- | --- | --- | --- | --- | --- |
| **Counts** | 113 | 133 | NAN | 46 | 66 | NAN |
| **Age**<br>mean (sd) | 18.401<br>(5.232) | 17.540 (5.300) | 0.099 | 18.592<br>(4.985) | 18.342<br>(5.377) | 0.739 |
| **Sex**<br>female % | 49% | 56% | 0.158 | 54.8% | 45.2% | 0.185 |
| **IQ**<br>mean (sd) | 71.420<br>(13.060) | 111.393<br>(12.518) | <0.001 | 67.237<br>(11.302) | 74.251<br>(13.582) | 0.005 ** |
| **White matter CVR**<br>mean (sd) | 0.029<br>(0.013) | 0.027(0.014) | 0.097 | 0.027<br>(0.013) | 0.030<br>(0.013) | 0.105 |
| **Grey matter CVR**<br>mean (sd) | 0.059<br>(0.021) | 0.060 (0.023) | 0.515 | 0.055<br>(0.021) | 0.061<br>(0.022) | 0.058 |
| **Scanner type**<br>HUG % | 68.5% | 64.6% | 0.404 | 73.8% | 64.3% | 0.158 |
| **Congenital heart defect**<br>female % | 37 (54%) | NaN | NaN | 17 (53%) | 20 (55%) | 0.9 |
| **Anxiety disorder (%)** | 54 (48%) | NaN | NaN | 28 (61%) | 26 (39%) | 0.025 ** |
| **Attention deficit hyperactivity disorder (%)** | 56 (50%) | NaN | NaN | 23 (50%) | 32 (49%) | 0.875 |
| **Mood disorder (%)** | 27 (24%) | NaN | NaN | 15 (33%) | 12 (18%) | 0.079 |
| **Schizophrenia spectrum disorders (%)** | 11 (10%) | NaN | NaN | 10 (22%) | 0 (0%) | <0.001 ** |
| **More than one psychiatric comorbidity (%)** | 48 (56%) | NaN | NaN | 26 (67%) | 22 (49%) | 0.001 ** |
| **Second generation antipsychotic**<br>(History of ever taking) | 53 (33.5%)<br>42<br>unknown | NaN | NaN | 43 | 10 | <0.001<br>** |
| **Serotonin reuptake inhibitor**<br>(History of ever taking) | 138 (85.7%)<br>39<br>unknown | NaN | NaN | 68 | 70 | 0.039 * |

### MRI acquisitions and parameters

The structural images were obtained with a T1-weighted sequence comprising 192 slices, with a volumetric resolution of 0.86 × 0.86 × 1.1 mm^3^ (repetition time = 2500 ms, echo time = 3 ms, field of view = 23.5 cm^2^, flip angle = 8°, acquisition matrix = 256 × 256, slice thickness = 1.1 mm, phase encoding right > left, no fat suppression). BOLD resting state fMRI scans were obtained using a T2-weighted sequence (200 frames, acquisition matrix = 94 × 128, field of view = 96 × 128, voxel size = 1.84 × 1.84 × 3.2 mm^3^, 38 axial slices, slice thickness = 3.2 mm, repetition time = 2400 ms, echo time = 30 ms, flip angle = 85°, phase encoding anterior > posterior, descending sequential ordering, GRAPPA acceleration mode with a factor for parallel imaging = 2). During resting-state scans, participants were instructed to fixate on a white cross on the screen, remain awake and let their mind wander (Zöller, Padula et al. 2018).

A total of 448 scans were acquired and were first included in the pool of analysis. However, 34 scans were excluded during quality control due to having a rotational or translational movement of above 3mm during the scan. Four scans had a brain scan with the vertex of the brain out of the field of view and therefore were excluded. One scan was excluded because the subject fell asleep during the scan. Finally, 409 scans were retained for the calculation of the CVR.

### Preprocessing and CVR map creation

The data analysis was carried out using Statistical Parametric Mapping (SPM) software (SPM12 Wellcome Trust Centre for Neuroimaging, London, UK; http://www.fil.ion.ucl.ac.uk/spm)) and custom MATLAB scripts (MathWorks, Natick, MA). The preprocessing steps for the BOLD image series included motion correction, smoothing with a Gaussian filter at a full-width-half-maximum (FWHM) of 6 mm, and linear detrending. Following the methodology of Liu et al. we obtained a proxy of the pressure of arterial CO2 changes based on intrinsic BOLD fluctuations, by filtering the BOLD time series within a 0.02–0.04 Hz frequency band (Liu et al., 2017). The average BOLD time course for this range was then computed considering a gray matter mask derived from a T1-weighted MPRAGE image, serving as the reference time course for gas fluctuations. Note that, because of the BOLD acquisition parameters, the cerebellum was removed from the field of view. Subsequently, a linear regression analysis was conducted using the reference time course as the independent variable and each voxel’s time course as the dependent variable. Motion parameters were also included as confounding factors. This analysis generated a CVR index for each voxel, resulting in a qualitative resting state CVR map for each participant at each visit. These maps were then warped into DARTEL space (Ashburner 2007) and further into the Montreal Neurological Institute space (Collins, Zijdenbos et al. 1998) for groupwise analysis.

### Mixed-effect model

To study the effect of age and clinical group (22q11 vs HC) on global CVR, the average white matter (WM) and gray matter (GM) CVR values for each subject were calculated using the WM and GM mask provided by the SPM segmentation separately. These values for the longitudinal sample were then used as dependent variables in two separate mixed-effect GLM models, one per tissue, with age, grouping and their interactions as independent variable, as well as sex as a covariate, using the publicly available toolbox (https://github.com/danizoeller/myMixedModelsTrajectories):

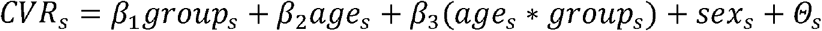

Where CVR_s_ is the average WM or GM CVR value of subject *s*, group_s_ is the group of subject s, age_s_ is the age of subject s, sex_s_ is the sex of subject s, and *Θ*_s_ is the random effect of the *sth* subject. We repeated the same analysis for a second set of mixed-effect models on a subset of our subjects, i.e. comparing PPS+ vs nonPPS groups.

Note that a mixed effect model was selected to account for multiple visits of individuals.

### PLS-C

To investigate the longitudinal trajectory of multivariate patterns of CVR, we applied a partial least squares correlation (PLS-C) (Krishnan, Williams et al. 2011). Similar to prior published work of this group(Delavari, Sandini et al. 2021, Forrer, Delavari et al. 2024), we added longitudinal age regressors on the behavior side of the myPLS toolbox, for which the implementation is publicly available (https://github.com/MIPLabCH/myPLS).

To briefly describe this analysis, the cross-covariance matrix (R) is computed between a set of design variables (Y) and the CVR maps per subject at each visit (X). The design side (Y) consists of variables encoding the binary variable of diagnosis (or psychosis susceptibility grouping); as well as the longitudinal age regressors and their interaction. These longitudinal age regressors are described as follows: The “mean_age” that encodes for each visit of the subject the average age of that subject across all visits. The “delta_age” assigns the difference between the chronological age of a subject at the time of the scan and the subject’s mean age across all visits. Of note, by design these two age regressors are orthogonalized and can be interpreted as the cross-sectional and longitudinal effect of age, respectively. We next devise additional regressors to calculate the interaction of group and age regressors to enable the model to show deviation of the developmental trajectories of one group with respect to the other. The covariance matrix (R = Y^T^X) is then computed and undergoes singular value decomposition (R = USV^T^) resulting in latent components (LC)s. These LCs are tested for significance by permutation testing (1000 permutations). The component was considered significant if the p-value was less than 0.01. This threshold was determined by multiple comparisons correction using the Bonferroni method, dividing the usual significance level of 0.05 by the number of components tested (5), resulting in a corrected significance level of 0.01. To check the stability of the results and account for the possibility of overfitting, we obtained bootstrap ratios obtained by performing bootstrapping with replacement (500 bootstrap samples). Details of the PLS-C are outlined in the supplemental materials. Bootstrap ratio scores were thresholded at an absolute value greater than 3, which corresponds to a 99% confidence interval not crossing zero and indicates a stable contribution from this variable to the LC (Krishnan, Williams et al. 2011).

## Results

A total of 409 longitudinal scans were included in the analysis. Among the 246 subjects (130 female), 113 were participants who carried the deletion (22q11 group – 46 PPS+) and the rest were the control group (HC). The table of demographic demonstrates the characteristic of the participants and tests regarding the possible differences within groups.

### Global GM and WM CVR

Average CVR of grey matter and white matter voxels were compared across groups over the age-span.

No group or age-interaction difference of average GM and WM CVR was detected when comparing 22q11.2DS versus HC nor when comparing across groups with and without psychosis vulnerability.

### Whole brain CVR (22q11 vs. HC)

The longitudinal voxel-wise CVR analysis comparing the age trajectories in deletion carriers and HC, using PLS-C, resulted in one significant component (p-value = 0.002) (Figure 2).

**Figure 1.**
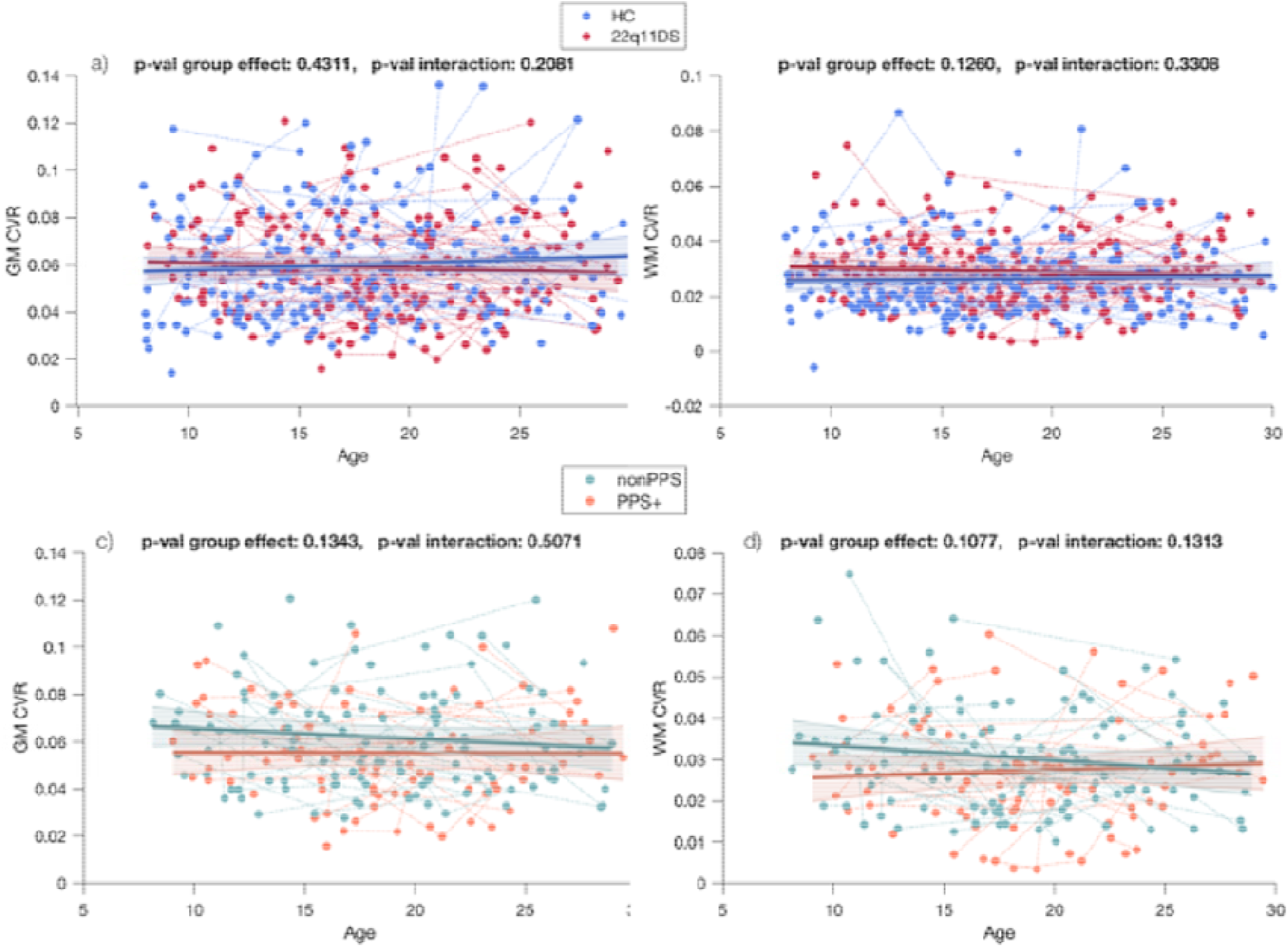
Comparison of global gray matter (GM) and white matter (WM) trajectories in groups. The significance of the effect of group (22q11 vs. HC or PPS+ vs. nonPPS) is demonstrated as “p-val group effect”. The significance of the effect of group (22q11 vs. HC or PPS+ vs. nonPPS) and age interaction is demonstrated as “p-value interaction “. Sex was included in all four models as a covariate. The group-level trajectory (solid line) is derived from mixed-effect model). The colored bands indicate the 95% confidence interval around the estimated group-level trajectory. Subjects’ repeated time-points are connected by dashed lines.

**Figure 2.**
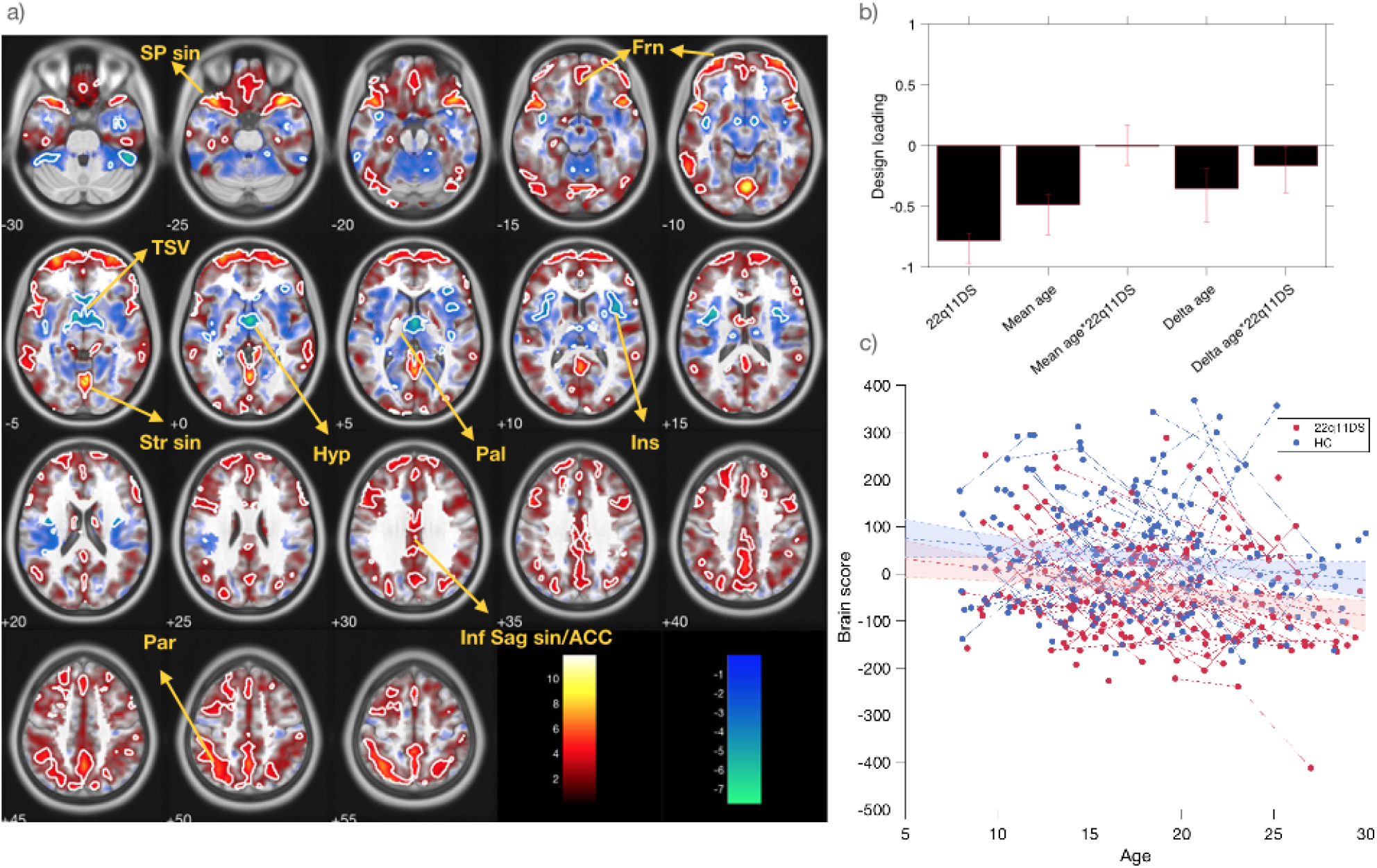
Multivariate voxel-wise pattern of CVR trajectories in 22q11.2DS versus HCs. Panel A) presents the spatial results of the component for which the colors indicate the effect size of the contribution each voxel to the component. The transparency of each voxel indicates the statistical strength of the contribution each voxel to the component. Clusters for which the strength is indicated statistically stable (as indicated by a high bootstrap ratio) are contoured with the white line– SP sin: Sphenoparietal Sinus; Str sin: Straight Sinus; Inf Sag sin: inferior sagittal sinus; ACC: Anterior cingulate cortex; Frn: Frontal; Par: parietal; Ins : Insula; Hyp : Hypothalamus; TSV : Thalamostriate veins

This component captured a steady age-effect as indicated by the stable age loadings as well as a prominent group-effect as indicated by the stable loading on 22q11.2DS diagnosis (Figure 2-b). As illustrated in figure 2-c, the group-age relationship is indicative of linear age relationship in both groups with a significant early offset in deletion carriers. Figure 2-a demonstrates the multivariate brain pattern associated with this age-diagnosis changes. Two sets of regions are revealed by this PLS-C. A first set of regions displayed in red demonstrate a steady decrease in CVR with lower values in participants with 22q11.2DS. This decrease is mainly observed within the major vasculature of the brain such as Sphenoparietal, Straight, and Inferior Sagittal sinuses as well as anterior cingulate cortex (ACC). Similarly, regions within the frontal and parietal lobe demonstrated a decreased CVR within the 22q11.2DS group. Conversely, within the Hypothalamus, Insula, and Thalamostriate veins, which are demonstrated in blue, CVR values increased with age while deletion carriers showed higher CVR values with regards to their HC counterparts. It is worth noting that both set of regions are revealed within a multivariate analysis. As such, participants with 22q11.2DS who demonstrate a decrease of CVR in red regions concurrently demonstrate an increase of CVR in blue regions.

### The whole brain CVR (PPS+ vs. nonPPS)

The PLS-C comparing the age relationship between PPS+ and nonPPS group revealed one significant component (p-value = 0.002) (Figure 3).

**Figure 3.**
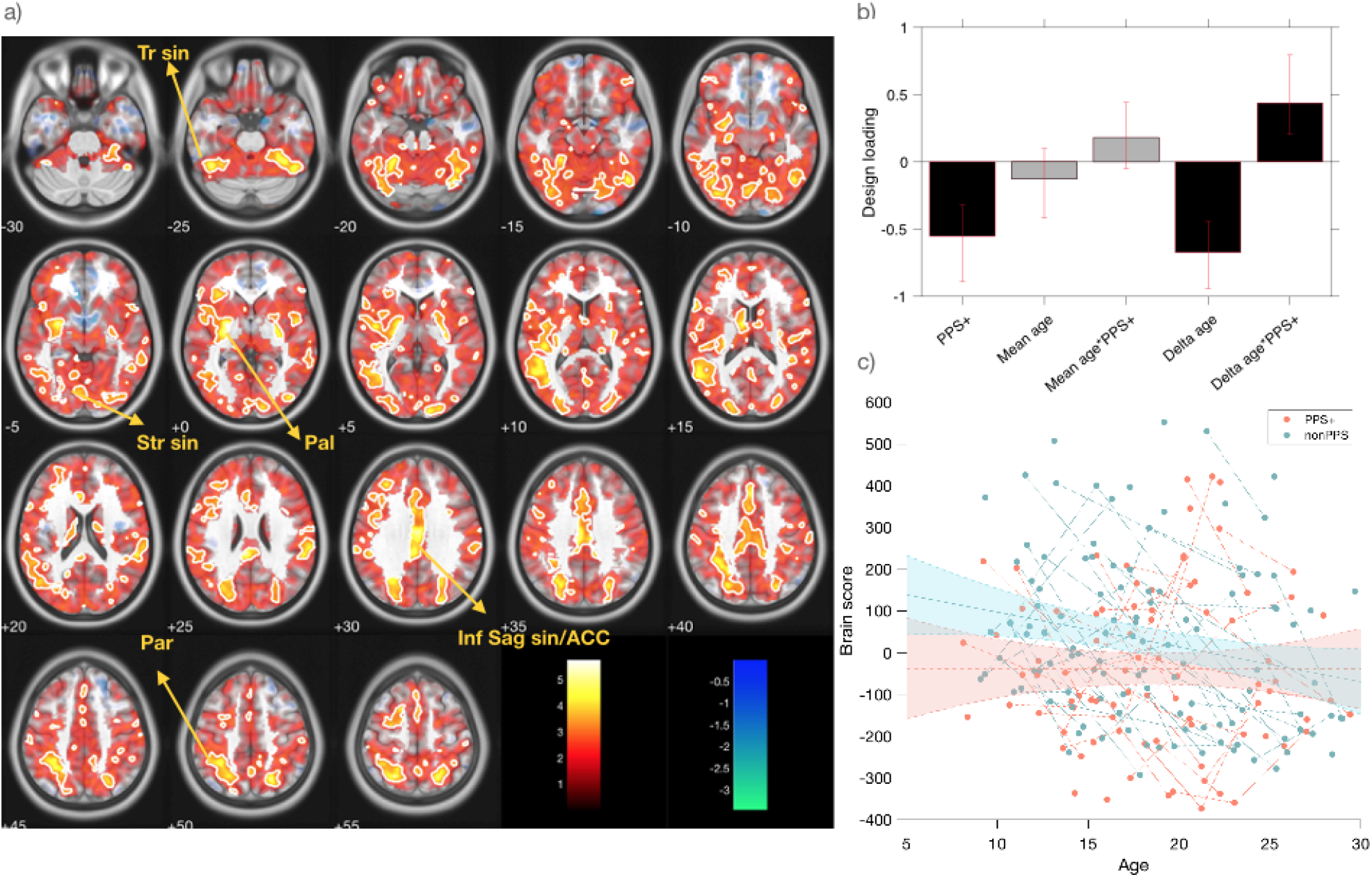
Multivariate voxel-wise pattern of CVR trajectories in PPS+ versus nonPPS. Panel A) presents the spatial results of the component for which the colors indicate the effect size of the contribution each voxel to the component. The transparency of each voxel indicates the statistical strength of the contribution each voxel to the component. Clusters for which the strength is indicated statistically stable (as indicated by a high bootstrap ratio) are contoured with the white line. _Str sin: Straight Sinus; Inf Sag sin : inferior sagittal sinus; ACC: Anterior cingulate cortex; Par: parietal; Tr sin : Transverse Sinus; Pal : Globus Pallidus

This component revealed a multivariate brain pattern mainly within Inferior Sagittal sinus, ACC Transverse sinus, Straight sinus, Globus Pallidus, and clusters within the parietal lobe (Figure 3-a). These clusters showed a prominent negative loading on the PPS+ group (Figure 3-b) as well as a positive age-interaction loading.

As illustrated in Figure3-c, this PSL-C revealed an early decrease of CVR within the above-mentioned regions in PPS+ group, whereas the nonPPS group reached this level of low CVR later in the aging process.

### Post-Hoc analysis investigating possible confounds

We performed 3 complementary analyses to investigate the effect of CHD in CVR, as well as two possible confounding variables namely antipsychotic medication intake and IQ which are significantly different across PPS+ and nonPPS groups. None of these PLS-Cs resulted in a component reaching the significance level. Nonetheless, we visualize and discuss the first component of each of these PLS-C in the supplementary materials.

## Discussion

To the best of our knowledge, this is the first study to examine CVR and in-vivo markers of cerebrovascular health in patients with 22q11.2DS. We conducted a multivariate analysis on a longitudinal cohort of 22q11.2DS to explore whole-brain CVR and its developmental alterations across deletion carriers, focusing particularly on a subpopulation with increased susceptibility to psychosis. This study provides initial evidence of an early hit to cerebrovascular health and its role in the pathophysiology of psychosis.

### CVR alterations during development

When investigating global WM and GM cerebrovascular reactivity, no age-related changes were detected. However, a voxel-wise assessment revealed anatomical regions where CVR varied with age. Little evidence is at hand regarding CVR alterations during development, with the majority of studies focusing on older populations (Liu, Li et al. 2017, Catchlove, Parrish et al. 2018, McKetton, Sobczyk et al. 2018, Peng, Chen et al. 2018, Pelizzari, Laganà et al. 2020, Solis-Barquero, Echeverria-Chasco et al. 2021, Ryman, Shaff et al. 2023). However, our results align with studies on the effects of aging on CVR, showing that while global WM and GM CVR remain stable over time, regional variability may be present across different brain areas in relation to age (McKetton, Sobczyk et al. 2018). Additionally, it has been demonstrated that the rate of vascular aging may differ across various brain regions, with one study showing that the temporal lobe experiences the fastest rate of longitudinal CVR decline in middle-aged individuals, followed by the parietal and frontal lobes (Peng, Chen et al. 2018). Our results demonstrate a steady decrease in CVR within frontal, parietal and striatal regions during development during adolescence and young adulthood.

As expected, the majority of clusters in the multivariate pattern showed a decrease of CVR with age. The decrease in CVR during aging is well documented (Catchlove, Parrish et al. 2018, McKetton, Sobczyk et al. 2018, Peng, Chen et al. 2018, Pelizzari, Laganà et al. 2020) and has been associated with normal loss of vascular wall elasticity and environmental factors such as inflammation. The decrease we observed during childhood and adolescence may as well be associated with the normal loss of elasticity in vasculature that already starts during childhood in central and peripheral blood vessels (Senzaki, Akagi et al. 2002). Remarkably, the multivariate nature of this study allowed for detection of a set of regions with a relative increase in CVR over age. This observation may be related to “steal” phenomena (Conklin, Fierstra et al. 2010), where lower CVR of certain regions flushes the blood volume into other regions with more CVR capacity. Such areas might be initially sub-performing compared to other regions (Arteaga, Strother et al. 2014). Alternatively, a reduction in CVR might be related to the development of a different hemodynamic delay in CVR (Stickland, Zvolanek et al. 2022), especially for regions characterized by bigger vessels. An interesting case is the increase of CVR over age in the hypothalamus. The hypothalamic paradoxical change in BOLD-derived CVR is reminiscent of studies demonstrating the inverse neurovascular coupling in the hypothalamus area due to neurons secreting vasopressin (Roy, Althammer et al. 2021). It is possible that hypothalamus demonstrates a contradictory CVR response of vasoconstriction to accommodate the specialized function of the region, similar to evidence in literature in nuclei within brainstem specialized for breathing, that paradoxically show vasoconstriction upon increase of CO2 (Wenzel, Hansen et al. 2020) to signal respiratory stimulation.

### Linking cerebrovascular health to psychosis susceptibility

Overall, our multivariate results showed that during childhood CVR is higher in clusters within the frontal and parietal lobe and major brain sinuses, and paradoxically lower within the hypothalamus and insular regions. Moreover, this normative CVR profile would become less present as participants grow. In addition to the age effect, we demonstrate that participants with 22q11.2DS show an early offset with regards to their CVR development, and are more similar to older HC individuals than their chronological age. Indeed, participants with 22q11.2DS may exhibit a defect in their CVR profile due to several interrelated factors. Individuals with 22q11.2DS are more susceptible to immunosuppression and thus multiple infections (Jawad, McDonald-McGinn et al. 2001). Moreover, metabolic disturbances such as obesity are more prevalent in the 22q11.2DS population (Voll, Boot et al. 2017). The chronic state of inflammation raised by obesity or multiple infections may eventually accelerate vascular aging and compromise CVR as suggested by the literature (Frosch, Yau et al. 2017, Spencer, Bell et al. 2018, Chiarelli, Villani et al. 2022).

Nevertheless, while findings indicate an early offset in CVR in deletion carriers, this pattern does not aggravate over time. Therefore, the causes presented at birth may be more plausible to contribute to this pattern. Among these causes, congenital factors inherent to 22q11.2DS, such as cardiovascular abnormalities and vascular anomalies are pivotal. Contrary to our initial hypothesis we did not find any alterations of the CVR in patients who had a history of congenital heart disease and those who did not. However, independent from presence of CHD (Unolt, Versacci et al. 2018), the deletion carriers are haploinsufficient for the claudin-5 gene (CLDN5) which leads to endothelial tight junction dysfunction and disruption of blood-brain barrier integrity which may contribute to the compromised CVR in 22q11.2DS population (Crockett, Kebir et al. 2024). Indeed, BBB impairment has been directly linked to CVR decline in animal models. In a recent study in rat brains, when the BBB was locally opened using focused ultrasound, attenuated CVR response, measured by the CO2 gas challenge, occurred in the affected region (Todd, Zhang et al. 2019). Therefore, our results regarding CVR reduction in 22q11.2DS may be in part linked to the established BBB impairment in this population.

Moreover, deletion carriers who eventually developed mild to moderate PPS+ demonstrated an early decrease in CVR within the ACC, frontal lobe, and the GB. In contrast, the group of patients without such susceptibility to psychosis (nonPPS) only exhibited this level of decreased CVR as a result of aging, by adulthood. This early reduction in CVR in these critical regions might thus play a crucial role in the later development of psychosis. Notably, the significant role of the ACC and frontal regions in schizophrenia, as well as in 22q11.2DS, has been well documented (Mattiaccio, Coman et al. 2018, Padula, Scariati et al. 2018). Additionally, the GB plays a key role in a critical circuit involved in psychosis, which includes the hippocampus, ventral striatum, and ventral tegmental area (VTA) (Figure 4). Briefly, increased excitatory inputs from the hippocampus to the ventral striatum decreases the inhibition of the ventral pallidum, resulting in heightened activity of VTA neurons projecting to the striatum (Egerton, Grace et al. 2020). This results in a downstream hyperdopaminergic state, which is believed to be a major underlying mechanism of psychotic symptoms (McCutcheon, Abi-Dargham et al. 2019).

**Figure 4.**
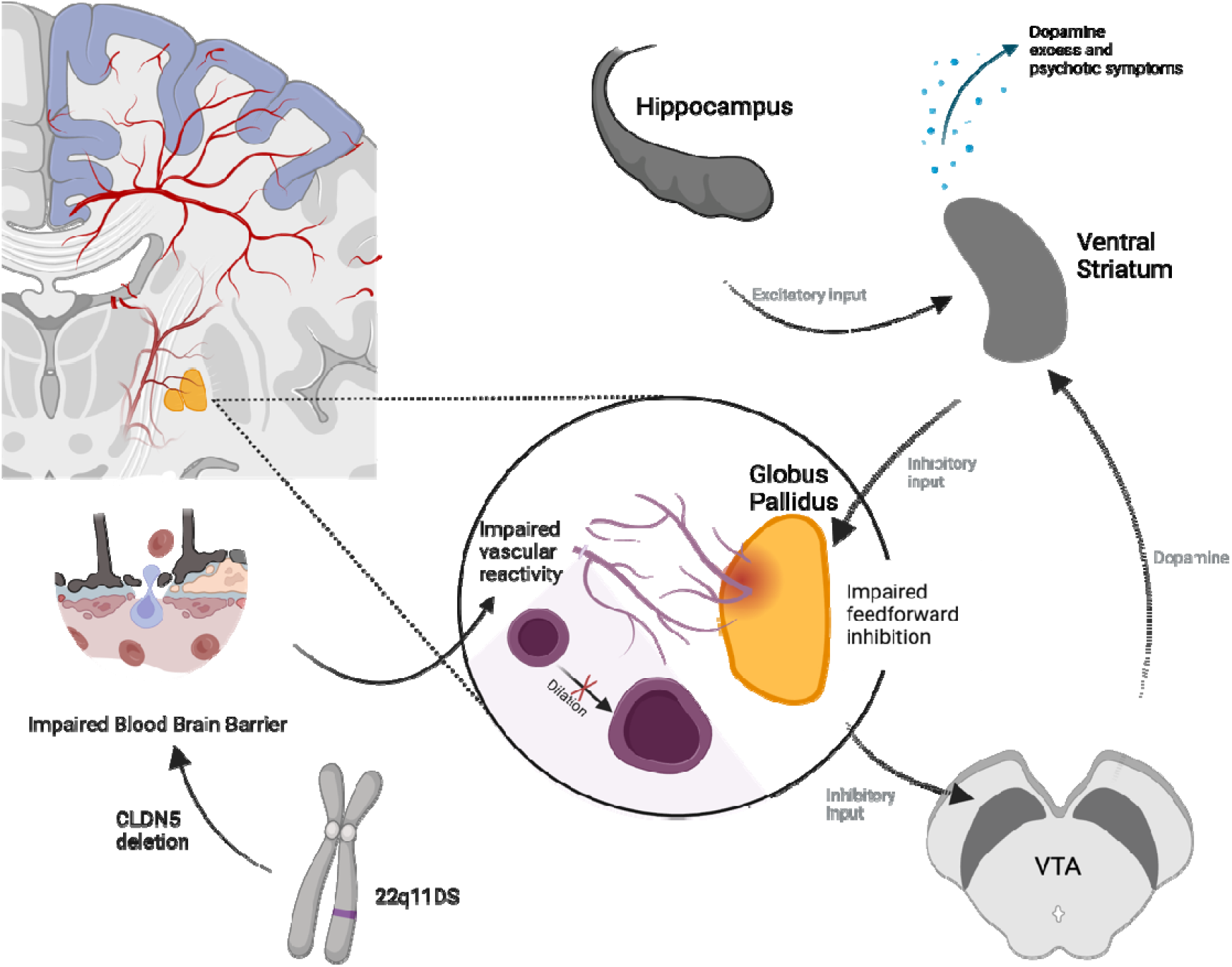
Putative mechanism through which cerebrovascular reactivity may increase the risk for development of psychosis. Cerebrovascular reactivity may represent blood-brain-barrier impairment, which in turn results in abnormal development and malfunctioning of susceptible regions. Among those regions, the globus pallidus plays a key role in the feed-forward inhibition of sub-cortical dopaminergic signaling.

Of note, the majority of deletion carriers first express psychotic symptoms during early adolescence (Schneider, Debbané et al. 2014). However, our results revealed the pattern of decreased CVR to be present already at childhood. Therefore, it is possible that early dysmaturation of the GB, due to abnormal CVR, may lead to a more pronounced lack of feedforward inhibition, making the population more susceptible to eventual hippocampal hyperactivity and the downstream hyperdopaminergic state that is documented during adolescent in this population (Delavari, Sandini et al. 2021, Mancini, Saleh et al. 2023). Therefore, the cerebrovascular health of the striatum may present an additional factor as an “early hit” that plays a role in the complex pathobiology of psychosis. This finding aligns well with the literature discussing the specific susceptibility of striatum vasculature due to particular anatomical features of this region. The striatum is served by the lenticulostriate arteries with narrow diameters, rendering them more vulnerable to vascular damage. This is significant as over 40% of strokes stem from severe blockages within the basal ganglia’s vessels (Feekes and Cassell 2006, Park 2016). Additionally, hippocampus and the striatum are shown to be more susceptible to BBB impairment within the cortex (Senzaki, Akagi et al. 2002), with recent evidence showing BBB impairment within the hippocampus is more transient, but persists within the striatum. Notably, a recent study (Zachariou, Pappas et al. 2024) showed that lower CVR within the striatum was associated with markers of BBB impairment. Overall, this evidence suggests the critical role of the vascular health and BBB integrity, especially within the striatum, for normal development. In light of these findings, our findings regarding a decreased CVR in young individuals with 22q11.2DS could remark the early BBB impairment in these individuals.

### Methodological considerations and future perspectives

The results presented in this study must be interpreted within the context of several methodological considerations. Firstly, we used resting state BOLD recordings for the calculation of CVR. Even though this method has been verified and utilized in several studies (Pinto, Bright et al. 2020), the gold-standard for measuring CVR remains methods incorporating a gas challenge. Indeed, performing such specialized sequences in children with a rare genetic deletion remains a big challenge, therefore we leveraged the flexibility of resting-state data to benefit from a long-term longitudinal study with a big sample size. This means that our CVR maps are only qualitative, not quantitative measurements.

Second, disentangling the effect of medication intake and PPS+ presents a significant challenge as inevitably, the majority of the PPS+ group have a history of taking antipsychotic medications. In order to better understand the effects of medication, we performed a secondary analysis using history of taking second generation antipsychotics as the grouping contrast. This PLS-C did not lead to a component reaching the significance level. Albeit the pattern of the first component in this non-significant PLS-C was similar to the one of the PPS+ vs nonPPS PLS-C (Fig S3). However, as antipsychotic medication is shown to improve blood flow in the striatum (Goozée, Handley et al. 2014), it is unexpected that the reduced CVR is derived by the medication intake. A more detailed investigation regarding the effect of medication is available in the Supplementary materials. Future studies with information regarding onset of medication intake as well as duration and dosage of medication are needed to further delineate the role of medication intake in CVR.

Additionally, increased CVR in 22q11.2DS within certain brain regions was contrary to our initial hypothesis. Indeed, the multivariate nature of our analysis is not allowing for deducting conclusions on individual regions and results should be considered as a pattern. Therefore, the increase of CVR within regions of the brain happens concurrent to the decrease of CVR to other regions. Future studies incorporating a lagged-CVR methodology may be helpful to further elucidate these results and whether this observation could be attributed to the steal phenomena.

In short, we showed a substantially early decrease of CVR values in a group of deletion carriers with eventual development of positive psychotic symptoms. We demonstrate this decrease is present years before the onset of symptoms and is most prominent within the GB, a key region for inhibiting dopaminergic cascades. Hence, we provide the first evidence linking cerebrovascular health in childhood to psychosis susceptibility in 22q11.2DS. Based on the trajectories of the alterations, decreased CVR may present an early biomarker of psychosis susceptibility. We propose a model in which cerebrovascular integrity must be considered among the first hits in the pathophysiology of psychosis, possibly through early damage to the striatum caused by neuroinflammation and mediated by BBB impairment. Further research should be conducted to disentangle the possible relationship of BBB impairment and CVR within the critical brain regions in young individuals with 22q11.2DS. These studies could elaborate whether this biomarker is also a potential treatment target for restoring BBB function as a preventive measure for psychosis.

## Supporting information

Supplemental Matrials

## Acknowledgments

This work has been supported by the Swiss National Science Foundation (SNSF) (Grant No. 320030_212476 [to SE]) and a National Centre of Competence in Research (NCCR) SYNAPSY (Grant No. 51NF40-158776 [to SE]). The authors report no biomedical financial interests or potential conflicts of interest. We would like to extend our appreciation to the families and the participants who took part in our cohort. We thank Greta Moltrasio and Caren Latrèche for the coordination of the project and the MRI operators at Campus Biotech, namely Roberto Martuzzi, Loan Mattera, and Nathalie Philippe.

## Notes

### Competing Interest Statement

The authors have declared no competing interest.

