## Supplemental Matrials for "Aberrant cerebrovascular reactivity presents as an early biomarker of psychosis susceptibility in patients with 22q11.2DS"

**Details of included variables in PLS-C**

In the PLS-C design variables, we begin by incorporating a binary diagnostic variable. Depending on the analysis, this variable separates either HCs from patients with 22q11DS or PPS(+) from non-PPS individuals. Considering the study's longitudinal nature, we devised age-related variables as outlined below to capture both overall cohort aging effects and individual age-related changes.

Firstly, we use mean-age, defined as the average age of each subject across all study visits. Secondly, delta-age represents the difference between a subject's age at a specific visit and their mean-age. As detailed in our previous publication, the orthogonalization of mean-age and delta-age ensures they correspond to cross-sectional and longitudinal aging effects, respectively.

The matrix U contains a left singular vector for each latent component, representing the saliences of seven design variables. These saliences indicate how each design variable contributes to the brain-design correlation. The diagonal matrix S holds the singular values for each latent component, quantifying the extent to which each component explains the correlation. The matrix V, containing the right singular vectors, represents the brain saliences, which express the contribution of each voxel to the correlation captured by the component.

**Results of the post-hoc analysis**

**CHD**

The PLS-C comparing the difference of CVR development in groups of deletion carriers with and without CHD resulted in first component with p-value=0.014.

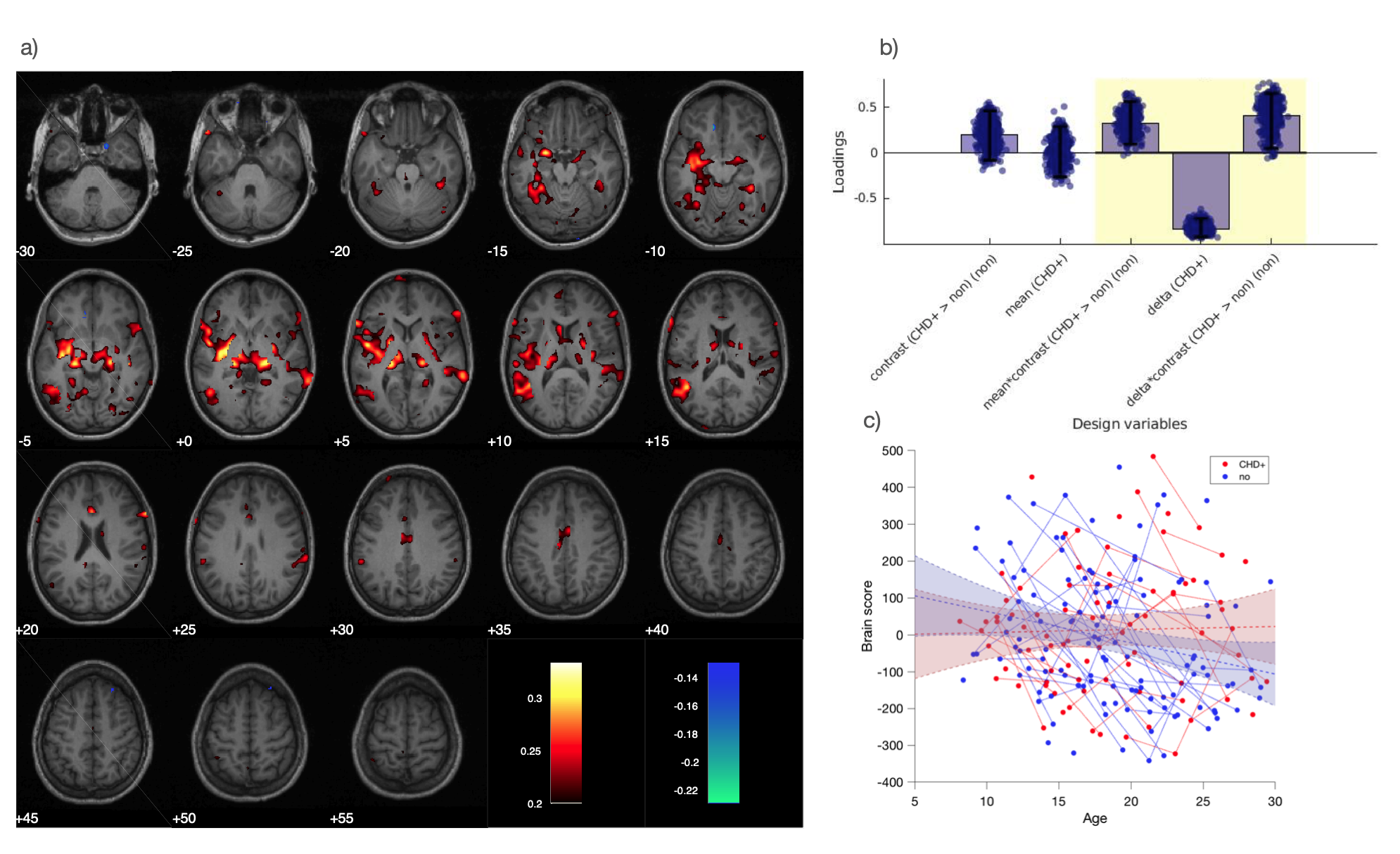

Figure S1

However, as depicted in Figure S1, this component captured mainly the effect of aging and the CHD grouping had minimal effect with bootstrap-ratios crossing 0 (Figure S1-b).

**IQ**

A similar component (p-value=0.011) with major age pattern was observed when comparing deletion carriers with IQ higher or lower than median IQ = 70. As depicted in figure S2, this PLS also showed minimal and unstable loading on IQ grouping (Figure S2-b).

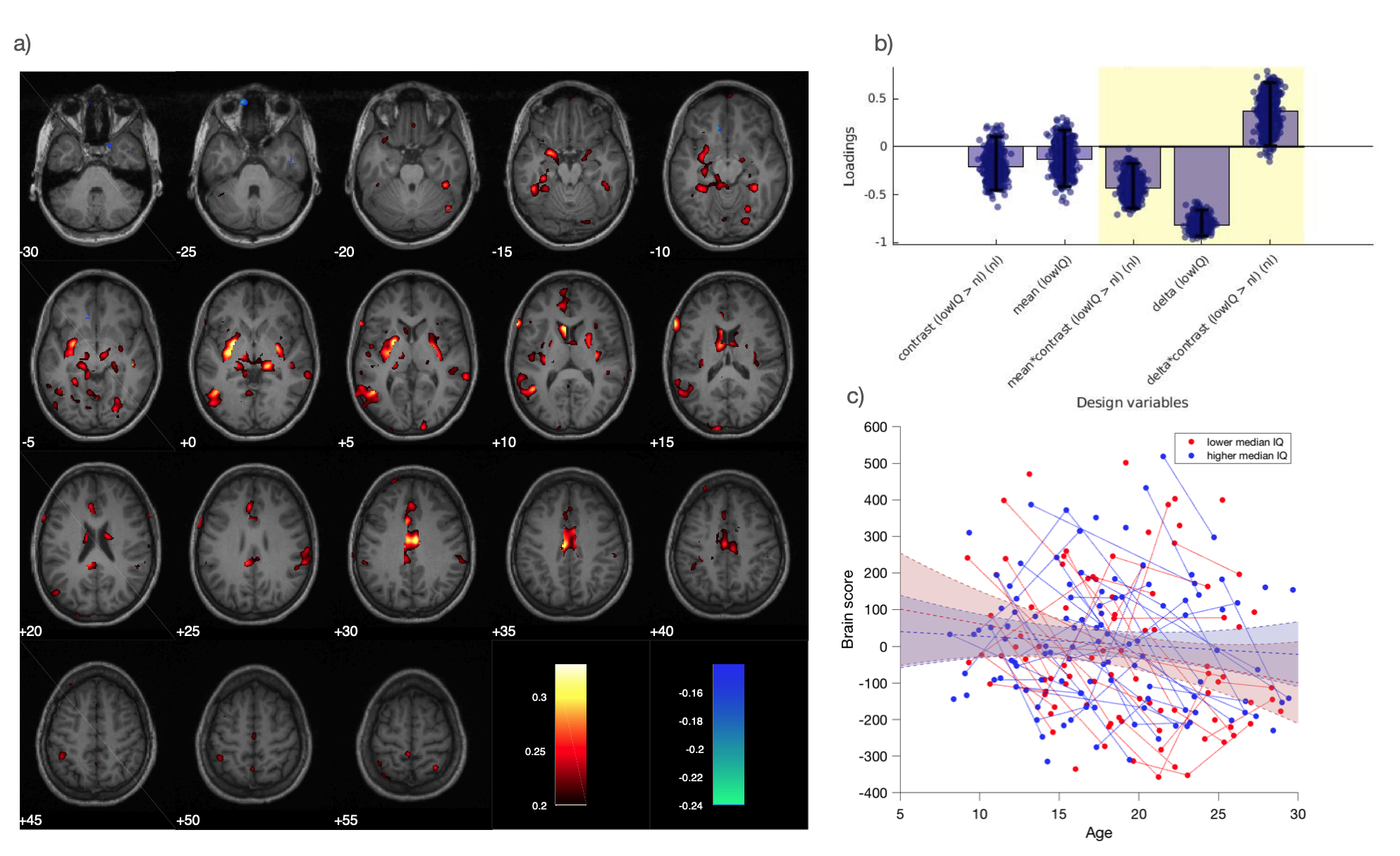

Figure S2

**Antipsychotics intake**
The PLS-C comparing deletion carriers who ever took second generation antipsychotic medication (Medication+) with those who did not (nonMedication), resulted in a first component with a p-value = 0.013.

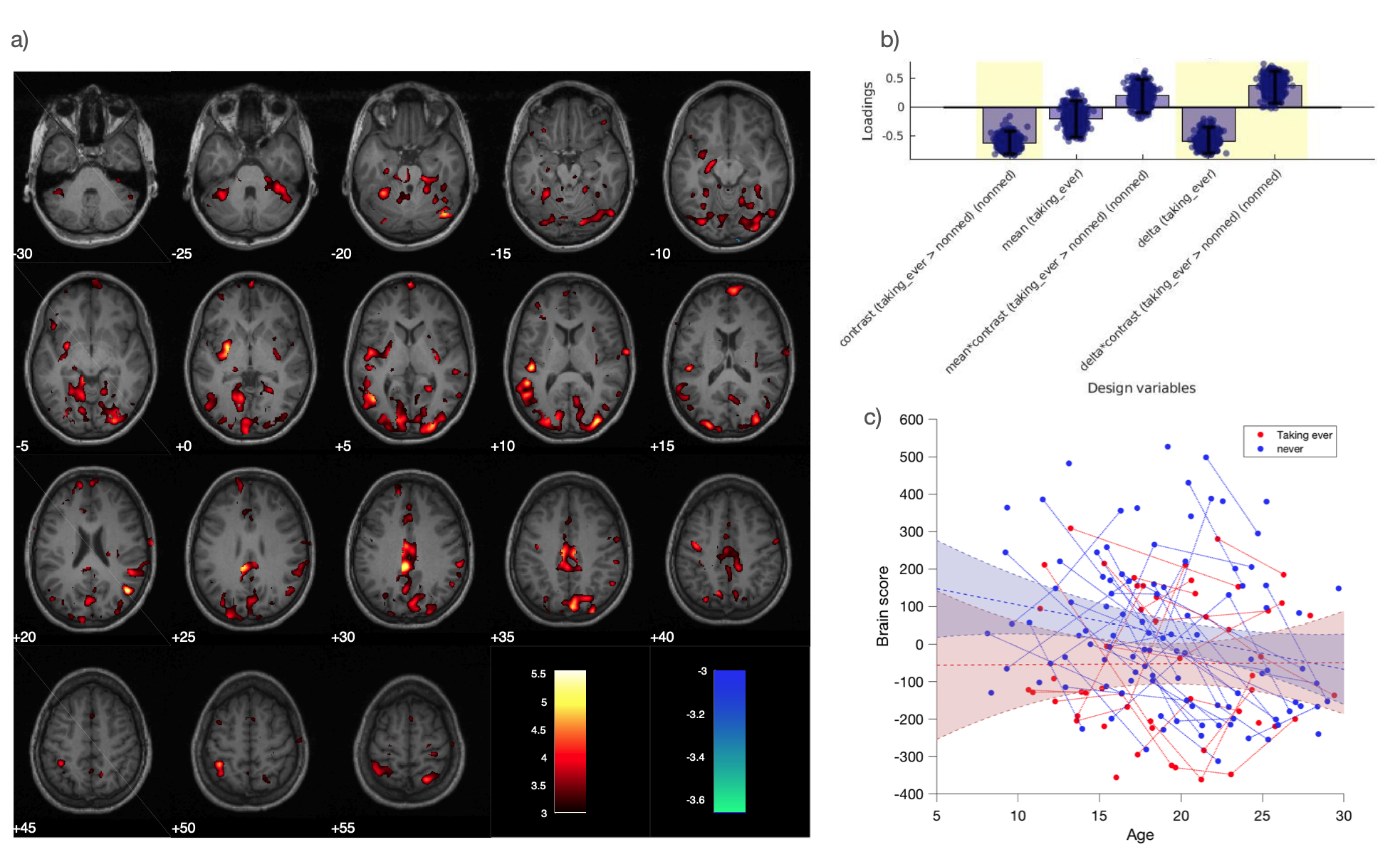

Figure S3

This component was very similar to the result of the PLS-C for PPS+ and nonPPS. These results are expected as inherently, there is substantial overlap between patients who take the medication and the PPS+ patients (Refer to Table-1 in the main manuscript). However, the group effect of the medication intake is smaller than the one of the PPS+.

Additionally, we performed a partial correlation analysis. We calculated the partial correlation of the brain-scores of the component demonstrating reduction in CVR, with both variables of PPS+/nonPPS and Medication+/ nonmedication. This analysis showed no significant partial correlation of the brain-score and the medication intake.

| Partial correlation with brain-score | Partial correlation co-efficient | P-value |
| --- | --- | --- |
| History of taking second generation | 0.0582 | 0.42 |
| PPS+ | 0.1616 | 0.023* |

We believe these results suggest the effect of belonging to the PPS+ group has a stronger contribution to the brain pattern observed in the PLS-C when compared to medication intake. Therefore, the results observed on medication intake may be explained by the overlap of the populations who are PPS+ and have a positive history for medication.
